# Aligning functional network constraint to evolutionary outcomes

**DOI:** 10.1101/278663

**Authors:** Katharina C. Wollenberg Valero

## Abstract

It is likely that there are constraints on how evolution can progress, and well-known evolutionary phenomena such as convergent evolution, rapid adaptation, and genic evolution would be difficult to explain under the absence of any such evolutionary constraint. One dimension of constraint results from a finite number of environmental conditions, and thus natural selection scenarios, leading to convergent phenotypes. This limits which genetic variants are adaptive, and consequently, constrains how variation is inherited across generations. Another, less explored dimension of evolution is functional constraint at the molecular level. Some widely accepted examples for this dimension of evolutionary constraint include genetic linkage, codon position, and architecture of developmental genetic pathways, that together constrain how evolution can shape genomes through limiting which mutations can increase fitness. Genomic architecture, which describes how all gene products interact, has been discussed to be another dimension of functional genetic constraint. This notion had been largely discredited by the modern synthesis, especially because macroevolution was not always found to be perfectly deterministic. But debates on whether evolutionary constraint stems mostly from environmental (extrinsic) or genetic (intrinsic) factors have mostly been held at the intellectual level using sporadic evidence. Quantifying the relative contributions of these different dimensions of constraint is, however, fundamentally important to understand the mechanistic basis of seemingly deterministic evolutionary outcomes. In some model organisms, genetic constraint has already been quantitatively explored. Forays into testing the relationship between genomic architecture and evolution included studies on protein evolutionary rate variation in essential versus nonessential genes, and observations that the number of protein interactions within a cell (gene pleiotropy) determines the fitness effect of mutations. In this contribution, existing evidence for functional genetic constraint as shaping evolutionary outcomes is reviewed and testable hypotheses are defined for functional genetic constraint influencing (i) convergent evolution, (ii) rapid adaptation, and genic adaptation. An analysis of the yeast interactome incorporating recently published data on its evolution, reveals new support for the existence of genomic architecture as a functional genetic dimension of evolutionary constraint. As functional genetic networks are becoming increasingly available, evolutionary biologists should strive to evaluate functional genetic network constraint, against variables describing complex phenotypes and environments, for better understanding commonly observed deterministic patterns of evolution in non-model organisms. This may help to quantify the extrinsic versus intrinsic dimensions of evolutionary constraint, and result in a better understanding of how fast, effectively, or deterministically organisms adapt.

**Glossary:**

Evolutionary constraint [1]: the phenomenon of evolution producing a finite number of genomic and associated phenotypic outcomes from a near infinite number of possible genetic variants.
Genetic constraint: The portion of evolutionary constraint which is determined at the level of genes or their gene products, for example codon constraint or developmental genetic pathways.
Functional network constraint: The portion of network constraint attributed to the structure or architecture of gene interactions that can be expressed in form of a network. Networks consist of nodes (genes) and edges (functional interactions between these genes).
Orthogenesis, Structuralism: The idea that properties inherent in organisms can direct evolution. Structuralism bases these properties on functional relationships of components that organisms are made of. Orthogenesis usually also implies that evolution is directed towards a goal. This view is not accepted within the modern synthesis of evolution.
Genic evolution: The phenomenon of different evolutionary outcomes being the outcome of independent mutation and selection events in different genes. For example, the occurrence of convergent evolution in diverging populations, both of which are caused by evolution in distinct genes.
Rapid adaptation: The phenomenon of adaptive change in allele frequencies of a population to natural selection, taking place within just a few generations.
Convergent evolution/convergence: Similar phenotypes evolving from similar selective pressure. May be (but doesn’t have to be) caused at the genomic level through genomic re-use of the same genes or alleles, which is also called parallel genetic evolution or genomic re-use.
Gene dispensability: a variable to measure gene essentiality. The less dispensable a gene is for organismal growth and function, the more essential it is.
Pleiotropy and cost of complexity: Gene products with many functional interactions with other gene products are constrained to accumulate less nonsynonymous mutations, because this would negatively affect the phenotype in many ways. Consequently, more complex genome organisation leads to higher constraint.
Gene expression level CAI: The amount of mRNA produced by each gene in regular somatic cells. CAI (Codon Adaptation Index) is used as a substitute variable in this paper, and is derived from codon use bias in yeast that correlates with mRNA levels.
Omega ω: the ratio of nonsynonymous to synonymous substitutions dN/dS. It is assumed that dS remains constant, and dN is used as a measure for directional evolution.
Gamma γ: A score developed for quantifying or predicting events of rewiring functional connections between network nodes over the course of evolution. Developed on the example of five species of yeasts.
Neighborhood connectivity: A network statistic used to describe the structure of a functional genetic network. Describes the number of connections of all neighbors of each node. Highest values expected in intermediately located nodes
Betweenness centrality: A network statistic used to describe the structure of a functional genetic network, describing how a node lies within paths between other nodes. Nodes with many paths progressing through them may be important in transmitting information. Highest values expected in central nodes.
Average shortest path length: A network statistic used to describe the structure of a functional genetic network. Shortest distance between a node and other nodes. Highest values expected in peripheral nodes of a network.

## Genetic constraint and evolutionary outcomes

Understanding the nature of the genetic dimension to evolutionary constraint is crucial to understanding adaptation and repeatedly observed outcomes of evolution such as, for example, convergent evolution [2]. Guyer (1922) already pointed out a possible constraint of molecular level processes on selection, on the example of convergently evolved bird colour patterns being generated by the same enzymes [3]. Genetic constraint is here meant to be similar to the notion of structuralism (constraints on form), and distinguished from orthogenesis which describes the notion of a goal-directed evolution, that can even be nonadaptive, and stems from inherent organismal properties. Evolutionary constraint is here defined as the phenomenon of evolution producing a finite number of genomic and associated phenotypic outcomes from a near infinite number of possible genetic variants. Convergence means that similar phenotypes arise in phylogenetically distant lineages in response to similar environments [2]. Traditionally, convergence has been studied in non-model organisms, and with a focus on adaptive modification (e.g., [4]). But more recently, phenotypic convergence has been traced back to resulting from identical genotypic variants (called “genomic re-use”, revised in [5]). These can either arise as new parallel mutations, or from parallel selection of the same alleles from standing genetic variation [5] such as in the independent selection of body armor in the ectodysplasin locus of stickleback fish [6]. More tentative evidence for convergence at the molecular level, comes from the recent findings of genomic re-use in generating convergent adaptations. For example, two poison frog species with a most recent common ancestor estimated at 159 mya, evolved parallel changes in the same gene related to skin toxin transport [7]. A recent study of Yudin et al. [8] found several independent instances of parallel functional genomic adaptation to temperature in a range of extant and extinct mammals inhabiting extreme cold environments. Such genomic re-use causing convergence in distantly related lineages may indicate that constraint at the genomic level is important to generate convergent evolution. However, it has been argued that this may not be a universal phenomenon, as convergent phenotypic adaptations may alternatively be produced by different genes. They may also be exaptations, where a similar allele evolved due to ancestrally different selective pressures with a subsequent change of function [9]. At present it is unclear whether genomic re-use is a common basis for evolutionary constraint, and if so, how often it can be expected to occur against a backdrop of a myriad of possible mutated allele combinations and recombinations that exist in any given genome. Recent studies, including ours, have provided insights that an interplay between environmental (extrinsic) and organismal (intrinsic) constraints may generate convergent phenotypes [10,11], with intrinsic constraint being identified as a causative factor for speciation [12], but not many comprehensive evaluations of the effect size of genetic constraint on evolution have taken place yet.

Another puzzling outcome of evolution is rapid adaptation within diversifying populations as a rapid response to selection such as anthropogenic pollution [13], which can occur even in the presence of gene flow [13,14]. The speed of such adaptation and the simultaneous occurrence of both convergence (possibly due to genomic re-use) and divergence within the same genome has been dubbed the “genic theory of speciation” [14] and was shown to occur in *Timema* stick insects [15] (Table 1 H5). Further evidence for simultaneous convergence and divergence, combined with genomic re-use, was found in the extensively studied Caribbean lizards of the genus *Anolis* [16]. *In one such example, divergent genetic populations of Anolis cybotes* in the Dominican Republic have evolved convergent phenotypic, ecological, reproductive, physiological, and parallel genomic adaptations to high elevations on three separate mountain chains [17–20]. Certain components of the phenotype in this lizard diverged along the same axes of the phenotype that were modified during more ancestral stages of the *Anolis* radiation, while other components of the phenotype convergently adapted to elevation using previously unmodified phenotype components [18]. Genomic signatures of this adaptation to climatic clines were found in genes with functions known to be involved in climate adaptation across metazoans [20,21], indicating evolutionary re-use of the same genetic pathways to similar selection pressures across populations. Other recent research on *Anolis* lizards suggests that such simultaneous convergence and divergence can evolve over the course of just a few generations [22–25]. The generally rapid nature of such adaptations was a surprising common feature that emerged from the study of *Anolis* lizards [16,26]. Climate adaptation in *Anolis* can happen within even just one generation, as shown by genomic signatures of selection in *Anolis carolinensis* in response to a polar vortex descending on invasive populations in Texas [27]. Again, these phenomena suggest that the variants available to mutation and selection are constrained at the genomic level, indicating that genetic constraint is important for understanding outcomes of evolution, such as its speed, and upon which genes selection acts.

**Table 1.** Hypotheses relating network constraint to evolutionary outcomes and results of hypothesis assessment using a node classification scheme in yeast.

| Evolutionary outcome | Hypothesis No. | Hypothesis (H) | Alternative Hypothesis (HA) | Results in this paper following assessment with hierarchical node classification scheme. |
| --- | --- | --- | --- | --- |
| Speed of evolution | 1 | Indispensable or essential genes are more constrained and evolve slowly [77]. | Functionally important and thus functionally constrained genes evolve slowly, independent of dispensability [43].<br>Highly expressed genes evolve slowest [38,39]. | **HA:** Functionally most constrained genes (H-nodes) have the lowest substitution ratios of all categories, and are most highly expressed, but have lower scores of evolutionary rewiring than P and I-nodes. |
| Speed of evolution | 2 | Central nodes have highest number of edges, evolve very slowly because any change will lead to maladaptive pleiotropic effects - causes balancing selection through cost of complexity.<br>[5],[51],[40],[50] | Intermediate nodes evolve fastest as their higher number of edges allows for evolution through rewiring ([60,61], this contribution). | **HA:** Nodes with highest number of edges are intermediate to the network, evolve fast (high $\omega$) and have a high score of rewiring ($\gamma$), indicating that the substitution rate of these genes may be associated with evolutionary rewiring events. |
| Speed of evolution | 3 | Nodes with a low number of edges evolve fastest due to higher degrees of freedom which allows for genetic adaptations minimising pleiotropic effects.<br>[78], [5] | --- | **H:** Peripheral nodes evolve fast (high $\omega$) and have a high score of rewiring ($\gamma$), indicating that the substitution rate of these genes may be associated with evolutionary rewiring events. |
| Convergent evolution | 4 | Nodes with a low number of edges should be the prime target of convergent evolution. Pleiotropic negative effects are expected to be low, and mutations in them can maximise adaptation [5]. | Peripheral nodes have the highest degrees of freedom and thus divergence is more likely than convergence in them. Convergent evolution should instead be favored in nodes that allow for genetic variance, while having reduced degrees of freedom (I-nodes) (this contribution). | **HA:** 21 out of 26 nodes with convergent evolution demonstrated in yeasts were classified as I- nodes by DFA, and five as P nodes. $\omega$ and CAI were similar to I-nodes, but none of these 26 nodes showed evidence of evolutionary rewiring. |
| Genic evolution | 5 | Adaptations can be characterized (either causative or correlative for the speciation process) by any number of divergent genes within the genome, whereas other genes are not associated with adaptation. [14]. | Only the complete phenotype is selected, the genic component is less important [35]. | **H:** Different clusters of functionally similar nodes experience either higher, lower than expected, or neutral rates of evolution across five species of yeast [69]. Causation or correlation to speciation process not testable with data. |

During the recent decades, network thinking has emerged as a powerful approach to better understand biological realities [28]. Networks can lead to improved understanding of systems through their emergent properties [29] because systems might operate in ways that are unpredictable by only looking at their parts. Functional interactions between many genes in a cell can be modelled as gene interaction networks [30] which have been used to explain disease profiles and the functional genomic basis of cellular fates [31,32]. The network concept might also have deep implications in evolutionary biology. Futuyma [33] cited Schluter [34], noting that correlations between genes could reduce the degrees of freedom on which selection can operate. Many studies have shown the non-independence of genes from one another, be it through physical linkage, phylogenetic relationship, or functional interaction (Figure 1). Mayr [35] stated that “coadapted” genes are a result of natural selection, being brought together to form a “balanced system”, but ruled out that such gene complexes would be of any interest to evolutionary biology, as ultimately only the complete phenotype is selected [35], p.184ff (Table 1 H5A). While it is still unclear, and debated [36], whether genetic network structure is an outcome of evolution by natural selection, it could have implications for the understanding of selection from another, poorly explored perspective: genetic networks may serve as a mechanistic basis for genetic evolutionary constraint, and thereby could contribute to constraining and thus directing the evolution of genes contained within this structure, and the phenotype components associated with it [28]. The present contribution synthesizes empirical studies on genetic networks as being predictive for outcomes of evolution through genetic constraint, and explores new avenues for future research. I propose approaches to inferring the link between genetic constraint and three different evolutionary outcomes, and provide exemplary applications testing for them: **1)** convergent phenotypes based on parallel genomic evolution, **2)** the simultaneous occurrence of convergence and divergence within a genome, resulting in genic adaptation, and **3)** the existence of rapid adaptation.

**Figure 1.**
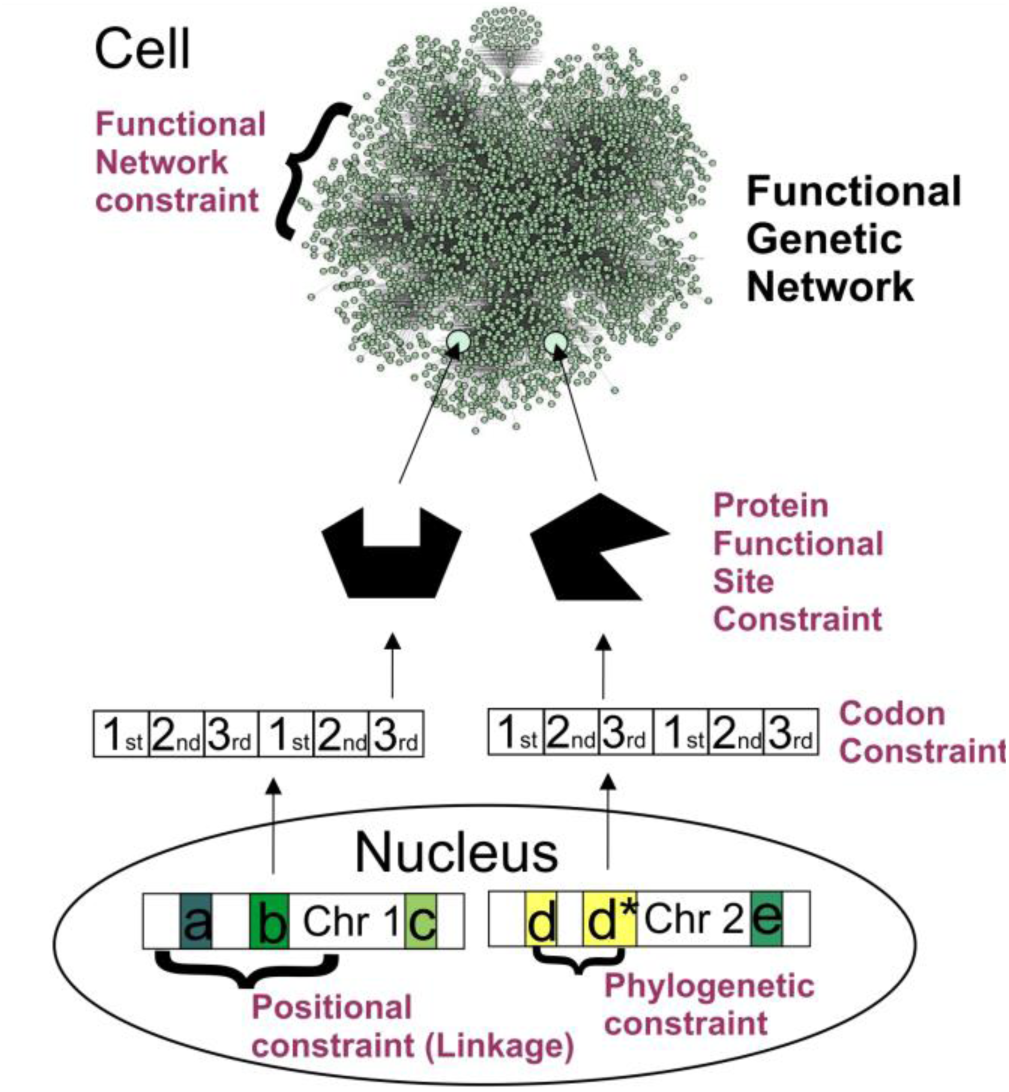
Examples for genetic constraint. On the DNA level, positional linkage determines recombination frequency, and gene ontogeny comprises a functional constraint (such as in the hemoglobin multigene family), that may be overcome by functional exaptation. Codon constraint determines how mutations accumulate within coding regions, since first codon position mutations change amino acid sequence of a protein, and have a higher likelihood to be deleterious than second and third codon positions. Codons within protein functional sites are constrained by protein function. Proteins and noncoding regulatory molecules such as RNAs form functional genetic networks (depicted is the functional network for genes involved in the human hypoxia response). Here, it is proposed that functional genetic network structure, as well as node position within the functional network, also pose a constraint to how beneficial mutations can accumulate.

## Genetic constraint through gene functional importance

The functional importance of a gene has long been thought to cause evolutionary constraint at the molecular level: protein-coding genes that are indispensable for the organism should be highly conserved and thus, be constrained through evolution, as most nonsynonymous mutations would be detrimental to protein function and thus most likely be non-adaptive. Consequently, these genes should have a lower rate of molecular evolution. Such genes have formerly been identified through their “**dispensability**”, which describes how essential they are for organismal function (Table 1 H1). The conserved *hox* genes causing vertebrate segmentation during embryonic development are commonly used as example for this hypothesis.

Zhang and Yang [37] reviewed evidence from empirical studies, and contrary to the expectation, found that across different model organisms including invertebrates and vertebrates, essential genes are not evolving more slowly than nonessential genes. Instead, highly expressed genes seem have lower rates of protein evolution (dubbed the “E-R anticorrelation” [38,39], Table 1 H1A), which some authors relate to translational selection on amino acids with different metabolic cost [38]. But perhaps functional importance needs to be defined differently than via gene essentiality or dispensability, and expression level may just be another correlative variable with another cause -- In *Saccharomyces cerevisiae* (in following: yeast), which was used for many studies on protein evolutionary rate and functional importance, essential genes are required for organismal growth and performance under optimal environmental conditions. A gene that renders an organism nonfunctional may thus predominantly be active in genetic pathways related to development and growth. However, in a complex organism such as a vertebrate, also the genes that are essential for organismal viability and reproduction are of high functional importance, and potentially could be under evolutionary constraint [40], such as in the example of genes coding for eye color determining mating success in *Drosophila melanogaster* [41]. A high proportion of the human genome has also been found to be under selective constraint in other mammals, indicating that gene dispensability is not a binary variable [42]. As reviewed by Zhang and Yang [37], Wilson et al. [43] suggested that evolutionary rate may instead be determined by **functional importance** and **functional constraint**. If functional importance measured as (negative) gene dispensability does not predict variations and constraints of evolutionary rate, perhaps functionally more constrained genes are the ones evolving slowly (Table 1 H1A). Prior studies have attempted to identify functional constraint in terms of protein functional sites, called protein functional site constraint in Figure 1, and the Neutral Theory [44] already identified codon constraint as important for evolution. The fact that gene products interact with others within functional genetic networks, is a less explored dimension for functional importance, and explanation for functional constraint of evolutionary rate (Figure 1). For example, a study by Jeong and colleagues [45] found that genes with many functional interaction partners are more likely to also be essential, which, however, does not provide enough evidence to extrapolate directly from functional constraint to evolutionary outcome.

## Genetic constraint through pleiotropy

Functional genetic network structure has been shown to affect evolutionary outcomes through pleiotropy in yeast: genes that interact with many other gene products are thought to be involved in many cellular functions and have multiple (pleiotropic) effects on the phenotype [46,47]. Fitness effects of mutations in them are partitioned across several phenotypic components, increasing the likelihood of maladaptive effects, which means that often they are more conserved through evolution and evolve more slowly [46]. Furthermore, proteins with many interactants may be constrained in the evolution of their functional sites to instances of co-evolution with the interactant genes, in order to maintain their functionality (Table 1 H2)[48]. A corresponding model of evolutionary constraint on evolution through pleiotropy that was explicitly based on functional network node hierarchy within an interactome, was proposed by Pavlicev and Wagner [49]. They argued that for genetic adaptation in a target gene to happen, selection has to overcome the inertia generated through stabilizing selection of the genes functionally connected with the target [49]. The premise of this model is that any change in genotype-phenotype interaction represents a change in a developmental pathway and, due the position of a gene within a network, will have pleiotropic effects on the phenotype [49]. This pleiotropic effect was found to be small for most genes, but genes with large phenotypic effect also are more pleiotropic [40]. High pleiotropy is assumed to have a cost for adaptation, which was explained as nodes central to a network evolving slower [40,50]. This idea, dubbed the “cost of complexity” [51] would lead to faster evolution of organisms with less complex genomic architecture due to this constraint being relaxed [40], and to adaptive selection on standing genetic variation preferentially to occur in genes with low pleiotropic effects [5](Table 1 H3). With regards to evolutionary outcomes, pleiotropy was suggested to limit events of genomic co-evolution [48], genomic adaptation [5], and convergent evolution [5] in nodes central to a network (Table 1 H4). Consequently, the properties of nodes within a functional genetic network, may be informative to understand their evolutionary constraint. However, genes with the highest number of interactants, or according to [46–49,51], the highest degree of pleiotropy, are not necessarily the nodes central to the network as assumed by Wagner and colleagues, but are instead nodes with intermediate position within a network [52]. The number of edges of a node may, consequently, not be sufficient to disentangle the effects of network structure on evolutionary constraint since it only measures one of a network’s many properties. More comprehensive measures of genomic network architecture may be necessary than just the number of functional connections a gene has with others.

## Genetic constraint through functional network architecture

Gene interaction networks were found to evolve either faster or slower than comparable genes functioning without being connected to others [53–56][57], so that the overall network architecture or hierarchy of genes within the network is likely to contribute to the speed and mode of evolution, regulated through functional constraint of nodes within the network. Different nodes within such networks may play different roles in evolution, resulting from hierarchical node organization. Box 1 outlines a hypothetical scenario of functional genetic network architecture influencing evolutionary outcomes: When selection acts upon a population (for example, through a sudden change in climate), advantageous mutations will be selected from standing genetic variation (allele frequencies). Organisms possess a finite subset of biochemical pathways (underlying functional genetic networks) that are related to temperature homeostasis [21,58,59], and that align to a finite amount of selected phenotype components. The population must adapt to the newly arising selective pressure through selection of non-deleterious mutations in one of these subnetworks, but not of mutations in any other subnetwork, as these are unrelated to the stimulus / organismal fitness in response to it, and would therefore not result in adaptation. This constrains the number of mutations in the genome that selection will operate on in this case, and thus determines the evolutionary response through genetic constraint. Second, and of high importance for the general mechanism proposed herein, node hierarchy within these subnetworks poses an additional level of constraint: and this additional level reduces the “evolutionary search space” for potential beneficial variants. This can be illustrated through reducing network structure to distinct types of nodes, which I outline below: Network nodes which are functionally important for the operation of the network (hub nodes central to the network - in following **H-nodes**), should be less likely to already harbor significant genetic variation in first- or second-codon positions or regulatory regions because of their high functional constraint. Consequently, genetic variation, as well as fast adaptation to an environmental selective pressure, should both be more likely to occur within non-hub nodes (Table 1 H3) within the subnetwork. Nodes with highest number of edges are intermediately positioned within a network (intermediate nodes, in following **I-nodes**) and were shown to have weaker selective constraint [60,61] than centrally positioned nodes, as they have lower functional constraint than H-nodes. Consequently, they should evolve faster (Table 1, H2A). This assumption differs from the pleiotropy hypothesis, which places the highest functional constraint on these nodes (cf., Table 1 H2). However, because of their high cost of complexity, adaptation in I-nodes should be highly constrained in terms of which genes can adapt (depending on the nature of their functional interactions) and how (through changing the wiring pattern with other nodes). Because of pleiotropy, adaptations that do evolve in these nodes should have a larger phenotypic effect, which combined with the reduced possibilities for adaptation, increases the likelihood for convergent evolution in them (Table 1 H4A). Genes peripheral in the network (peripheral nodes, in following, **P-nodes**) have higher degrees of freedom due to the lowest degree of pleiotropy and should be able to accumulate genetic variation with least cost. As a consequence, the population should already harbor more genetic variation within these peripheral genes that selection can operate on. Change in such nodes however, due to lower pleiotropic interactions, would result in less phenotypic effect and thus they are less likely to promote large evolutionary changes. In such nodes, divergence is more likely to accumulate than convergence (Table 1 H4A, Box 1).

The expectation is thus that different node types will differ in standing genetic variation due to the different genetic constraints acting upon them. H-nodes will be very strongly constrained and only can accumulate little standing genetic variation, resulting in a low potential for selection to operate on. I-nodes will harbor sufficient standing genetic variation but be under high functional constraint, so that selection can only operate on a limited amount of variants that all have multiple phenotypic effects. In different organisms, the same variants can be selected quickly due to this reduced search space, which leads to parallel genomic evolution resulting in convergent phenotypes. P-nodes will be least constrained, allowing a lot of variation but less sweeping phenotypic effects due to lower pleiotropy. Selection can operate on multiple variants in these, selective advantages are more likely due to the lower pleiotropy in more genes, so selection will less likely lead to convergence. All three evolutionary outcomes can be explained with this mechanism of constraint through functional genetic network structure.

#### Box 1 Proposed testable relationship between functional genomic network architecture, structural node position, and evolutionary outcomes

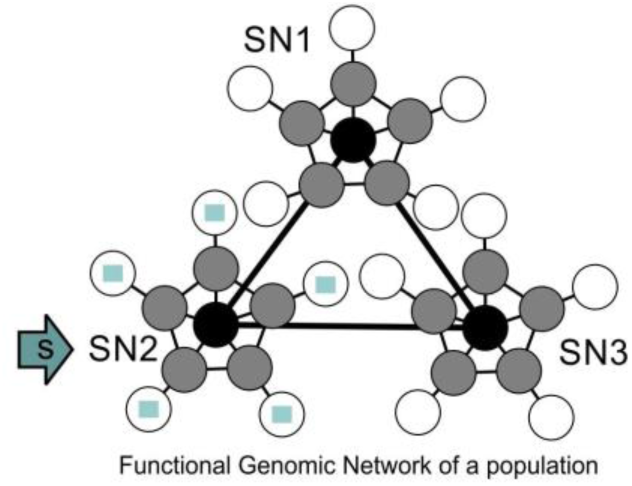

#### Constraint of network architecture

3 hypothetical subnetworks with 3 different tasks:

SN1: Structure development

SN2: Temperature homeostasis

SN3: Liver detoxification

#### Constraint of network node hierarchy

**Black** nodes are central for organismal function: not likely to accumulate non-synonymous mutations.

**Grey** nodes are functionally connected to many others: constrained in accumulating non-syn. mutations.

**White** nodes functionally connected to few others, least central for organismal function: most likely to accumulate non-synonymous mutations

#### Rapid adaptation

High standing genetic variation in white nodes allows for their adaptation. Selection being constrained to operate on those nodes that have accumulated such variation increases the speed of adaptation.

#### Convergent evolution

Adaptations to s are likely to happen in same functional subnetwork in different species / populations, since it is the one related to organismal response to the stimulus. The likelihood of convergent evolution is increased in grey nodes within that subnetwork, as they contain both genetic variance, and are constrained through a high number of network edges.

#### Genic evolution

Changing climate as a hypothetical selection pressure s is more likely to generate stress on SN2, than on SN1 and SN3 Selection is likely to operate on standing genetic variation, which is likely concentrated in white nodes. Adaptation to s is with high likelihood constrained to white nodes of SN2 (blue squares).

## Empirical hypotheses tests using data on yeast network evolution

A recent paper published by Schoenrock et al. [62] uses a data set of 4,179 protein-coding genes (sourced from [38,63]) to investigate the involvement of network structure in protein evolution. The data was obtained from five species of yeasts *(Saccharomyces cerevisiae, S. paradoxus, S. bayanus, S. kudriavzevii, and S. mikatae).* This study explicitly compares a quantitative variable related to network structure (computationally predicted re-wiring of nodes through evolution γ), with the protein evolutionary rate on nodes (substitution rate ω, measured as dN/dS). The authors find that the degree of rewiring of nodes across the phylogeny is only poorly associated with evolutionary sequence divergence, but nodes with very low evolutionary rate had high variability of rewiring scores, which indicates that changing gene interactions is an important mechanism how functionally constrained genes may evolve. While the study remained somewhat inconclusive about the influence of network structure and node rewiring on protein evolution, the data contained within it, combined with additional data, allow for a quick initial assessment of the three evolutionary outcomes outlined above (results are summarized in Table 1).

The most straightforward assessment can be done for outcome **(iii) Genic evolution** meaning differential evolution within the genome through selection on different subnetworks, as it is already discussed in the source paper [62]. Schoenrock et al. [62] could show that some nodes that are functionally similar experienced lower than expected levels of protein evolution, indicating purifying selection. Nodes that were evolving through fewer re-wiring events than expected, included functions related to phosphorylation, mitochondrial translation, response to pheromone, small GTPase mediated signal transduction, and transport. Nodes that were evolving among the five yeast species with higher than expected degrees of re-wiring, included the functions metabolic process, and various gene ontologies related to transcription and its regulation, as well as the regulation of transposition regulation. As indicated in Box 1, these results prove that evolutionary outcomes are different for functionally different subnetworks within an interactome. It might be worth noting that, as outlined above, none of these functions is particularly related to development but rather to maintaining organismal function, which is why they would be overlooked if conserved genes were only classified by the criterion of dispensability for colony growth. Gresham et al. [64] similarly showed that evolutionary constraint in experimentally evolved yeast populations over 200 generations is dependent on the type of selection (limiting Glucose or Phosphate vs. Sulphur), with convergence being an outcome of the system level organization of the respective metabolic pathway.

To assess evolutionary outcomes **(i) Rapid adaptation** and **(ii) Convergent evolution**, as well as to address the important factor of **gene expression** in shaping protein-coding gene evolution, the Schoenrock et al. [62] data set needs to be rearranged and expanded on. For this purpose, I obtained the data of [62] including yeast ORF ID, computationally predicted evolutionary PPI re-wiring score (γ), and substitution rate (ω). This data set was then integrated with data downloaded from Wall et al. [38] including ORF ID, and CAI (Codon Adaptation Index, a measure of RNA expression levels, based on [65]). With the goal to calculate a classifier that will aid in describing hierarchical node position within networks, common network statistical parameters were calculated from the yeast interactome in CYTOSCAPE v.3.6.0 [66] using the Network Analyzer function. Data for the matching node ORFs were appended to the data set, and variables with non-normal distribution were BoxCox transformed. The final data set contained 2209 ORFs. The parameters obtained from the yeast interactome were average shortest path length (maximal in peripheral nodes), neighborhood connectivity (maximal in nodes intermediate to the network), and betweenness centrality (maximal in nodes connecting subnetworks). Nodes with maximum values for each one of these three statistical parameters, and that were not overlapping with each other (1081 nodes, Figure 2), were each assigned to a category: P (peripheral nodes), I (intermediate nodes) and H (hub nodes). To assign node categories to the remaining nodes in the network that may be harder to visually allocate, a discriminant function analysis (DFA) was employed in STATISTICA (V13, Tibco).

**Figure 2.**
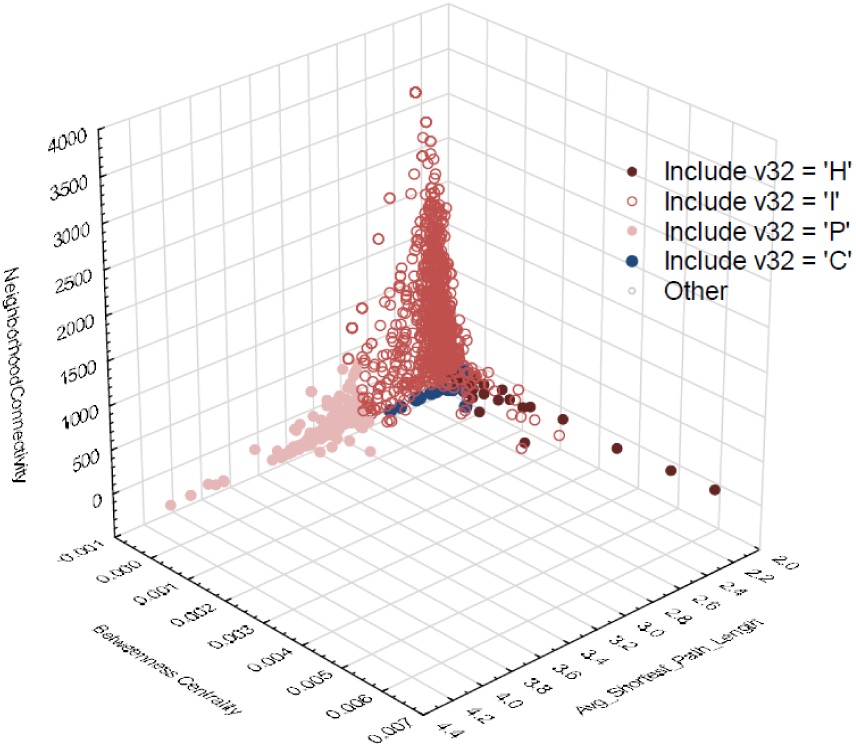
Distribution of yeast interactome nodes within network parameter space. (neighborhood connectivity, average shortest path length, and betweenness centrality). The top values for each axis are colored in shades of red (light, filled: P-nodes; light, open: I-nodes; dark, filled: H-nodes). Convergent evolution nodes are indicated in dark blue. These top values for each axis formed the basis to classify the remaining nodes based on discriminant function analysis.

All remaining nodes could with significant statistical support be associated to one of these three categories (Table 2). To explore the network position of nodes that have undergone convergent adaptation, ORF IDs that were demonstrated experimentally to show convergent genomic adaptation in independent experiments, strains, or species of yeasts (**C-nodes**) were identified from the literature ([67], [68], [69],[70],[64], [71], Table 3). Out of the 26 obtained C-nodes, 21 nodes were allocated by DFA to the I category, and five were allocated to the P-category.

**Table 2.** Discriminant function analysis summary to assign node categories H, I, P to nodes within dataset. Wilks’ Lambda: 0.0704 approx. F (6,2152)=992.780 p<0.001.

|  | Wilks<br>Lambda | Partial<br>Lambda | F-remove<br>2,1076 | p-value | Toler. | 1-Toler. (R-sqr.) |
| --- | --- | --- | --- | --- | --- | --- |
| Neighborhood<br>connectivity | 0.137 | 0.514 | 507.835 | <0.001 | 0.988 | 0.012 |
| Betweenness centrality | 0.105 | 0.673 | 261.039 | <0.001 | 0.994 | 0.006 |
| Average shortest path<br>length | 0.133 | 0.528 | 480.907 | <0.001 | 0.983 | 0.017 |

**Table 3.** List of yeast genes that were found to adapt to novel environments, and were additionally shown to evolve these adaptations convergently across populations or species of yeast. Node hierarchy categories after discriminant function analysis (DFA) are shown in the first column. P - peripheral nodes, I - intermediate nodes.

| DFA estimated Node hierarchy | Gene symbol | ORF ID | Reference |
| --- | --- | --- | --- |
| I | STE11 | YLR362W | Lang et al., 2013 |
| I | STE12 | YHR084W | Lang et al., 2013 |
| I | STE4 | YOR212W | Lang et al., 2013 |
| P | KRE6 | YPR159W | Lang et al., 2013 |
| I | SFL1 | YOR140W | Lang et al., 2013 |
| I | STE5 | YDR103W | Lang et al., 2013 |
| P | ANP1 | YEL036C | Lang et al., 2013 |
| I | GCN1 | YGL195W | Lang et al., 2013 |
| I | ERG5 | YMR015C | Gerstein et al., 2012 |
| P | ERG7 | YHR072W | Gerstein et al., 2012 |
| I | CNE1 | YAL058W | Lang et al., 2013 |
| I | GPB1 | YOR371C | Lang et al., 2013 |
| P | KEG1 | YFR042W | Lang et al., 2013 |
| I | KRE5 | YOR336W | Lang et al., 2013 |
| I | TOH1 | YJL171C | Lang et al., 2013 |
| P | SUL4 | YBR294W | Gresham et al 2008 |
| I | GAL3 | YDR009W | Hittinger et al., 2004; Stern, 2013 |
| I | GIN4 | YDR507C | Gresham et al 2008 |
| I | PDR1 | YGL013C | Anderson et al. 2003 |
| I | SGF73 | YGL066W | Gresham et al 2008 |
| I | SET4 | YJL105W | Lang et al., 2013 |
| I | SIR1 | YKR101W | Gresham et al 2008 |
| I | ACE2 | YLR131C | Lang et al., 2013 |
| I | GAS1 | YMR307W | Lang et al., 2013 |
| I | WHI2 | YOR043W | Lang et al., 2013 |
| I | CKA2 | YOR061W | Gresham et al 200 |

Figure 3 shows that DFA-allocated P, I, H, and C node categories significantly differed in the network statistics used to generate them. All network statistics significantly differed between node categories, as shown with Kruskal-Wallis tests: Average shortest path length: KW-H(3,2208) = 1220.5906, p = 0.0000; neighborhood connectivity, KW-H(3,2208) = 26.5707, p < 0.0001; betweenness centrality KW-H(3,2208) = 926.5707, p < 0.0001. It was then tested, how network statistical parameters relate to the evolutionary parameters ω, γ, and CAI. First, a general linear model was run with evolutionary parameters as dependent variables, and network parameters as predictor variables. All three network statistical parameters were found to significantly predict evolutionary outcomes (Table 4). All node categories have significantly different values for ω (KW- H(3,2204) = 20.1345, p = 0.0002), CAI (KW- H(3,2195) = 26.1472, p = 0.00001) and γ (KW- H(3,2195) = 36.7936, p = 0.00000), as shown by Kruskal-Wallis tests (Figure 4). With regards to hypothesis **(1) Rapid adaptation,** Figure 4 values (0.93 vs. 0.91), and lowest values were found in H nodes. This shows that nodes located less centrally in the network evolve faster than other nodes, but does not identify peripheral nodes as particularly fast-adapting. CAI increases towards the center of the network, with mRNA expression level being highest in hub nodes. Network node hierarchy may therefore be able to explain the E-R anticorrelation (gene expression levels being negatively correlated with evolutionary rate [37]): H-nodes connect various subnetworks with one another, and thus are likely to be involved in more diverse functions (which might be partitioned across different tissues, processes or life history phases), than nodes more peripheral in a network (Figure 4). Such common functions may require a high amount of product, which may translate into high levels of mRNA expression in these nodes. γ is highest in P and I-nodes, indicating that evolutionary rewiring events are more common in less central parts of the networks. I-nodes harbor the majority of edges within a network - genetically re-wiring these nodes could lead to rapid adaptation [72]. Centrality of H-nodes seems to reduce their adaptability while peripheral and intermediate nodes are less constrained to adapt, and this process may involve rewiring within the network. This demonstrates how functional constraint can explain evolutionary outcomes better than dispensability.

**Figure 3.**
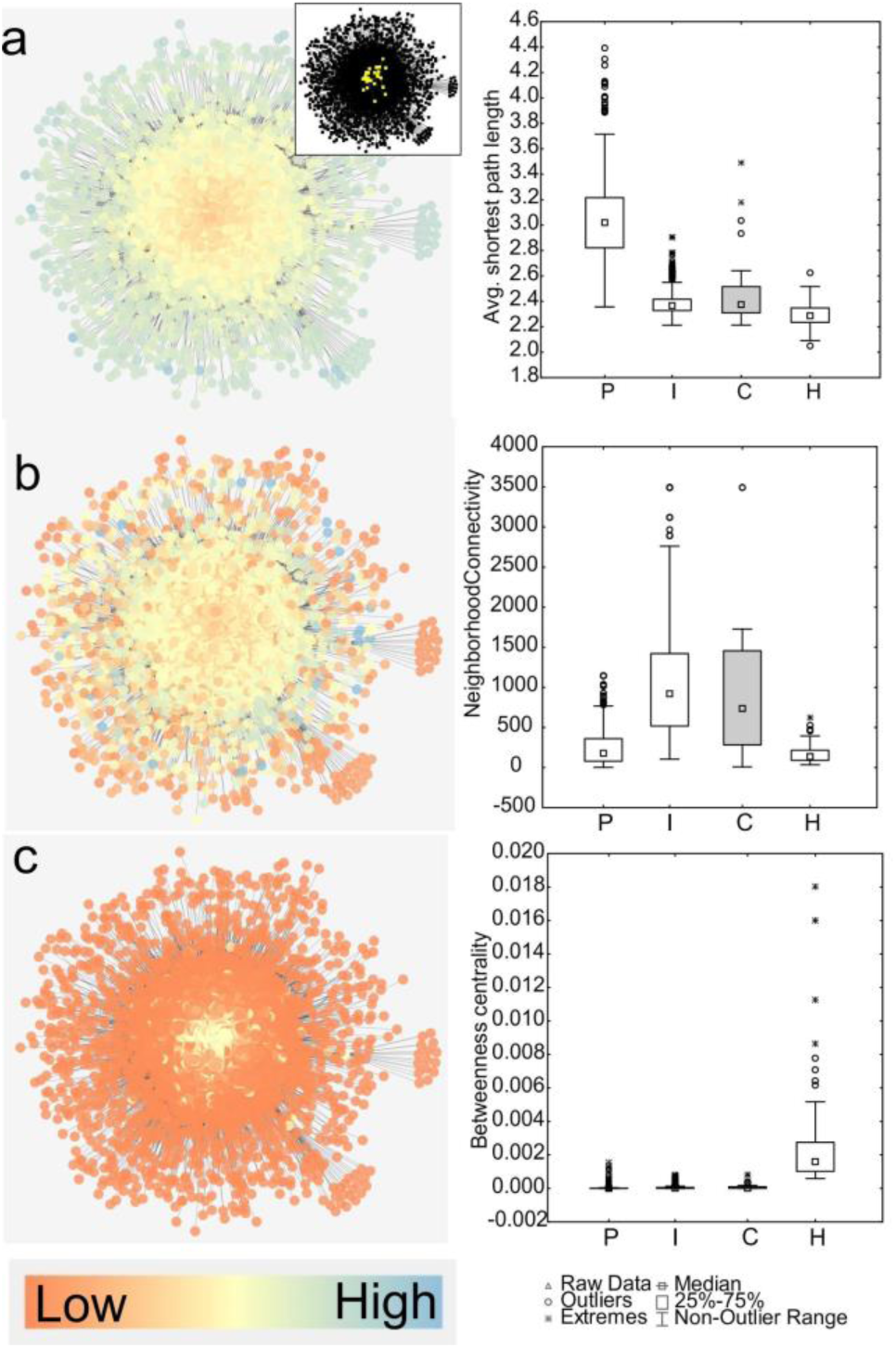
Visualization of node classification scheme in yeast interactome. Values of a) average shortest path length, b) neighborhood connectivity, and c) betweenness centrality within the yeast interactome (left panels), and values for the DFA-derived hierarchical node categories P, I and H, and for nodes known to be under convergent evolution in yeasts (C, N=18). Small inset network shows the location of convergently evolved genes (C-nodes) within the interactome (yellow, on black background).

**Figure 4.**
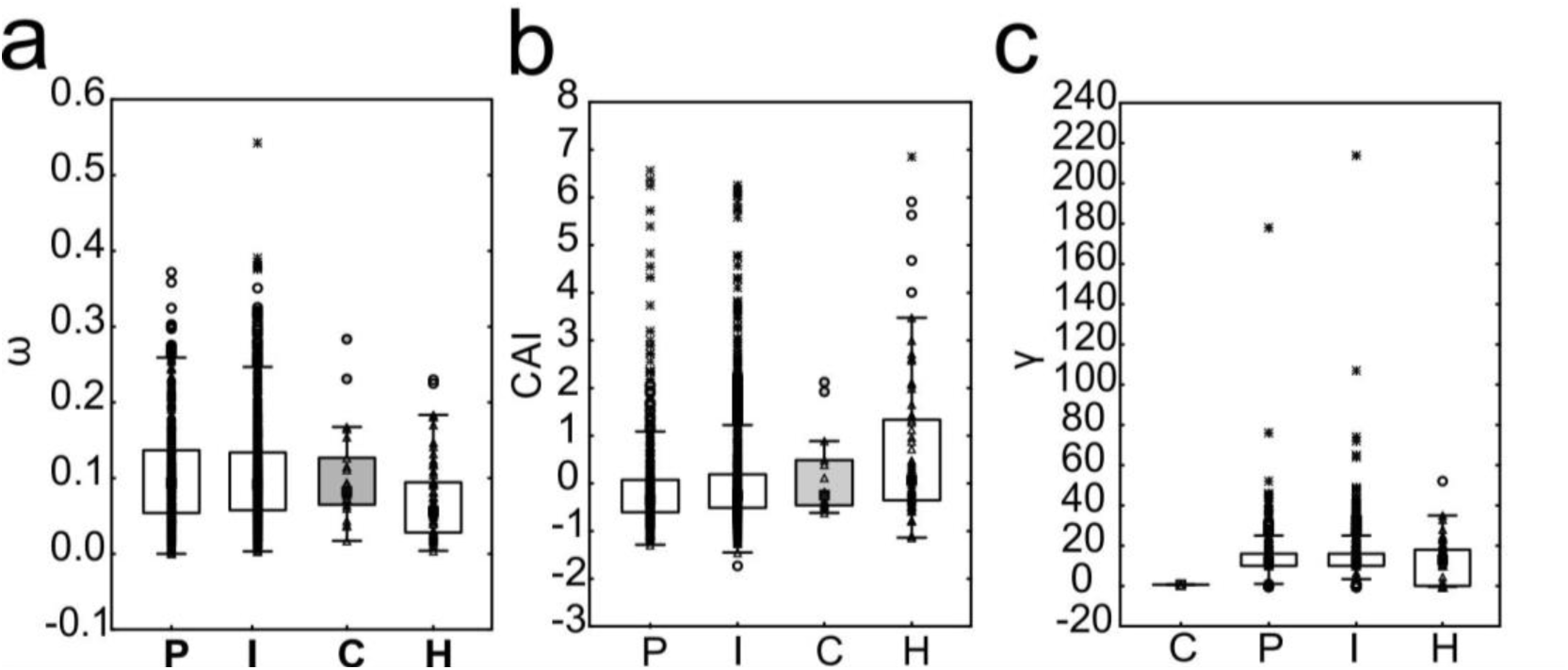
Relationship between hierarchical node structure of yeast interactome and evolutionary parameters. Node types are designated as peripheral (P), intermediate (I), or hub (H) based on discriminant function analysis, and nodes that were found to evolve convergently (C; N=22) in yeasts. Three evolutionary outcomes (a) substitution rate, (b) expression level, measured as CAI (Codon Adaptation Index), and (c) evolutionary rewiring score significantly differ among node categories (see text). C-node boxes are sorted by Median. Double red line: outliers above median not shown in figure but included in tests. Raw data points - triangles, circles - outliers, stars - extreme values, squares - Medians, boxes - 25-75% data, whiskers - non-outlier range.

**Table 4.** Multivariate Wilks tests of significance and powers for network parameters to explain protein evolutionary rate (ω), gene expression (Codon Adaptation Index CAI), and evolutionary rewiring between species of yeast (γ). All predictors were significant.

|  | Wilks' Lambda | F | Effect df | Error df | p | Observed power (alpha) |
| --- | --- | --- | --- | --- | --- | --- |
| Intercept | 0.317 | 1569.597 | 3 | 2188 | <0.001 | 1.000 |
| Neighborhood connectivity | 0.924 | 59.892 | 3 | 2188 | <0.001 | 1.000 |
| Betweenness centrality | 0.995 | 3.931 | 3 | 2188 | 0.008 | 0.832 |
| Average shortest path length | 0.961 | 29.553 | 3 | 2188 | <0.001 | 1.000 |

Hypothesis **(2) Convergent adaptation through network constraint** is also tentatively supported through Figures 3 and 4. C-nodes are most similar to I-nodes, showing that convergently adapting genes have similar evolutionary rates, expression levels, and degrees, as nodes that are located intermediately in the interactome. This supports the idea that nodes with highest number of edges and intermediate network position are constrained to adapt and thus increase the likelihood for convergent evolution. Gresham et al. [64], from which five C-nodes were obtained, also showed that convergent evolution is related to system level organization of the respective metabolic pathway. In summary, these results show a clear relationship between network node structure and evolutionary outcomes. An overview of all results linking network parameters to evolutionary outcomes is provided in Table 1.

## From yeast to non-model organisms

Metagenomic resequencing of every 500 generations within a 60,000 generation *E. coli* long term evolution experiment [73] revealed that certain genes accumulated beneficial mutations through selection significantly more often than expected by chance, and were very often affected by parallel adaptation [73]. These results, together with the incidences of parallel genomic adaptations reviewed herein, demonstrate that the above described relationship between network structure and convergent evolution may be expandable to organisms other than yeasts [5]. Apart from the quick assessment performed in this contribution, the influence of network structure in shaping evolutionary outcomes in more complex organisms than yeast such as vertebrates still needs to be comprehensively tested. As demonstrated above in the yeast example, the impending advent of large-scale functional genomic networks for many new species makes it possible to convert functional genomic network structure into variables describing hierarchical node position within the network. Future tests relating evolution to genomic constraint could include node architecture, and revolve around (1) Comparing standing genetic variation to network node position (while considering the effect of demography, selective sweeps, genetic drift, bottlenecks, and other levels of extrinsic constraint); (2) Testing whether similar subnetworks/node hierarchies adapt to same selection pressure in different organisms. (3) Comparing the speed of realized adaptation to a mutation/selection expectation, without considering network constraint. The potential benefits on better understanding genetic constraint leading to deterministic evolution may be wide ranging: in humans, the use of functional interaction networks is omnipresent in genomic and transcriptomic study of cancer data, and recently, calls have been made for evolutionary methods to be applied to cancer problems [74]. A recent study demonstrates how the early progression of pancreatic cancer is defined through evolutionary constraints resulting from following one of three tumor suppressive pathways, and thus may be predictable [75]. Previous criticism against recognizing network constraint as evolutionary force has centered around the idea that this would disregard evolution through natural selection [76]. Instead of upholding such a dichotomy, I argue that the goal should instead be to quantify “background genetic constraint” through functional network structure, in order to better allocate the remaining variance to mutation and selection in directing rapid, convergent, and genic phenotypic evolution.

